# Lithium directs embryonic stem cell differentiation versus hemogenic endothelium

**DOI:** 10.1101/2020.04.07.031054

**Authors:** Hayk Mnatsakanyan, Manuel Salmeron-Sanchez, Patricia Rico

## Abstract

The discovery that the definitive hematopoietic stem cells (HSCs) derive from specialized regions of the endothelium, known as the *hemogenic endothelium* (HE), shed a good deal of light on HSC embryonic developmental processes. This knowledge opened up new possibilities for the design of new strategies to obtain HSCs *in vitro* from pluripotent stem cells (PSCs). Previous advances in this field have shown that the Wnt/β-catenin signaling pathway plays a key role in PSC-derived HSC formation. In this work, we identified lithium, a GSK3 inhibitor, as an element capable of stabilizing β-catenin and inducing ESC differentiation in the mesoderm lineage and subsequently in the HE, highly consistent with the role of Wnt agonists on ESC differentiation. ESCs treated with 10 mM lithium express CD31+, Sca-1+, Nkx2-5+ and Runx1+ cells characteristic of HE cells. The ability of lithium-treated ESCs to further derive into HSCs was confirmed after defined maturation, resulting in rounded cell aggregates positive for fetal and mature HSCs markers, confirming the endothelial to hematopoietic transition. Our results represent a novel strategy for generating HSC *in vitro* as a multipotent source of stem cells for blood and muscle disease therapies.

## Introduction

Hematopoietic stem cells (HSCs) are multipotent stem cells capable of self-renewal and differentiation. They are the main source of the precursors of all classes of blood cells^1^. Image tracing experiments have supported the idea that definitive HSCs arise from specialized vascular endothelial cells that acquire blood-forming potential (hemogenic endothelium, HE)^2–4^. HE specification responds to the need to develop a functional circulatory system to provide oxygen and nutrients to all types of tissue during embryogenesis^3^. HE and definitive HSC development from PSCs is driven by Wnt/β-catenin signaling, activated by either Wnt or Wnt agonists such as GSK3 inhibitors^2^Λ Previous studies on CHIR99021 have shown that this pharmacological GSK3 inhibitor is capable of stabilizing β-catenin and promoting its transcriptional activity^6^,^7^ and thus inducing HE formation. The activation of the Wnt pathway by inhibiting GSK3 using CHIR99021 in the first stages of ESC differentiation followed by stimulation of the vascular endothelial growth factor (VEGF) further induces the differentiation of PSCs to HE and subsequently to HSCs^8^. Lithium is another well-defined GSK3 inhibitor and has been used for the treatment of bipolar disorder for decades^9^. However, its inhibiting effect on GSK3 in ESC differentiation is somewhat controversial, as shown by previous studies that suggest that murine ESCs treated with lithium differentiate towards neural cells^10^ instead of mesodermal lineage.

Although considerable efforts have been made to direct PSC fate towards hematopoietic lineages, the efficient synthesis of HSC precursors needs a deeper understanding of the molecular mechanisms that control HSC fate^11^. HSCs are now widely used in clinical transplantations as a cancer therapy and for other blood or immune disorders^11^, although their use is restricted due to the present limited primary HSC sources and questions of donor/receptor compatibility^11^-^12^, so that alternative methods are needed to overcome these limitations. PSC-derived HSCs can thus be considered as a major breakthrough in this regard, even though the low efficiency in synthesizing hematopoietic cells from PSCs is one of the major hurdles to this approach^11^.Here we define the lithium-activated mechanisms on murine ESC self-renewal and differentiation, focusing on lithium-mediated inhibition of GSK3β activity and how this effect can modulate both β-catenin expression and nuclear translocation, driving ESC differentiation towards the HE cells. We demonstrate the ability of these lithium-treated ESCs to further derive into HSCs after defined maturation and confirm the endothelial-to-hematopoietic transition. Since β-catenin-mediated mechanisms for HSC development are conserved in human and mouse species^5,8,13^ our results demonstrate a novel strategy for generating efficiently HSCs *in vitro* as a multipotent source of stem cells to be used in a variety of blood and muscular biomedical applications.

## Materials and methods

### ESCs culture

D3 murine embryonic stem cells (ESCs, ATCC) were cultured in basal medium (BM: DMEM high glucose (Lonza) supplemented with 10% Knockout Serum Replacement (ThermoFisher), 1% Fetal Bovine Serum (Gibco), 1% 100X Nucleosides (Millipore), 1% L-Glutamine (Sigma-Aldrich), 1% Non-essential Amino Acids (Sigma-Aldrich), 1% 100X Penicillin/Streptomycin (P/S, ThermoFisher) and 10 mM 2-Mercaptoethanol (Gibco)) at 37 °C in a humidified incubator containing 5% CO2, and 1,000 U/ml ESGRO Leukemia inhibitory factor (LIF, Millipore) to inhibit ESC differentiation. Before seeding, culture dishes were coated with 0.2% gelatin (Sigma-Aldrich). Lithium chloride (Sigma-Aldrich) was used as Li^+^ source for all the experiments.

### ESC viability and proliferation in the presence of different lithium concentrations

The cells were cultured on gelatin-coated plates to study the role of Li^+^ in ESC proliferation and viability at 10,000 cells/cm^2^ in BM supplemented with either different Li^+^ concentrations or 1,000 U/ml of LIF for 3 days. After 1 and 3 days, the ESCs were lysed by Tris/Triton X100/EDTA (10 mMTris pH 8, 0.5% Triton X100, 1 mM EDTA) buffer, and proliferation was calculatedf by quantifying total DNA concentration by a Quant-iT PicoGreen dsDNA Assay Kit (ThermoFisher).

### Hemogenic endothelium differentiation experiment

In order to assess Li^+^-mediated differentiation of ESCs into HE, the ESCs were cultured at 10,000 cells/cm^2^ on gelatin-coated plates for 6 days in BM and BM supplemented with 2 and 10 mM Li^+^. After the first 3 days of culture the cells were passaged and then sub-cultured for another 3 days.

After 6 days of culture, the HE cells developed from ESCs were characterized. HE cells were then matured to HSCs replacing BM with a maturation medium consisting of DMEM/F12 (Sigma-Aldrich) supplemented with N-2 (Thermofisher), B27 (Thermofisher) supplements, 1% L-Glutamine, 0.05% of BSA (Sigma-Aldrich) and 1% P/S, without added Li^+^. Cells were cultured in these maturation conditions for an additional 11 days.

### Immunofluorescence and stains

Cells were fixed with 4% formaldehyde for 20 minutes at room temperature (RT). The samples were then washed with TBS and blocked with TBS/Triton X100 0.1%/BSA 2% for 1 h at RT. Cells were incubated with primary antibodies overnight at 4° C (Table S1). After washing, samples were subsequently incubated with secondary antibodies (Table S1) for 1 h at RT. Hoechst (dilution 1:7,000, Sigma-Aldrich) was used for nuclear staining. Samples were mounted with 85% glycerol and imaged by a Nikon Eclipse i80 fluorescence microscope. The percentage of protein staining was quantified by image analysis on imageJ software.

### Western Blot experiments

Protein was extracted by RIPA buffer supplemented with protease inhibitor cocktail tablets (Roche). The proteins were separated in 10% SDS-PAGE with Mini-PROTEAN Electrophoresis System (Bio-Rad) and transferred to PVDF (GE-Healthcare) membranes by Trans-Blot Semi-Dry Transfer Cell (Bio-Rad). Membranes were blocked with 5% BSA (Sigma-Aldrich) or 5% skimmed milk (Sigma-Aldrich), in TBS/0.1% Tween 20 (Sigma-Aldrich) for 1 h and incubated with primary antibodies (Table S2) diluted in TBS, 0.1% Tween 20 and 3% BSA overnight at 4° C. Membranes were washed and incubated with HRP-linked secondary antibody (Table S2) for 1 h at RT. The chemiluminescence band was detected by ECL-Plus reactive (ThermoFisher). A Fujifilm Las-3000 imager device was used to visualize the protein bands.

### Chromatin immunoprecipitation

Chromatin immunoprecipitation was performed as previously described^14^. Briefly, cells were fixed for 15 min in 1% formaldehyde, and the reaction was stopped by adding 125 mM glycine (Sigma-Aldrich). After cell lysis, cell extracts were sonicated in a Bandelin Sonoplus Mini 20, using six pulses of 10 s (30% amplitude). Immunoprecipitation was performed using protein G Dynabeads (ThermoFisher) pre-adsorbed with anti β-Catenin antibody (ThermoFisher).

### qPCR experiments

Whole RNA was isolated by a Quick RNA Miniprep kit (ZYMO Research) and quantified by Q3000 micro volume spectrophotometer (Quawell). RNAs were retrotranscribed on a Maxima First Strand cDNA Synthesis Kit with Thermolabile dsDNase (ThermoFisher). Real-time qPCR was carried out using the PowerUp SYBR Master Mix (Thermofisher) and 7500 Fast Real-Time PCR System (ThermoFisher). Gene expression was quantified by the ΔΔCt method^15^. Sample values were normalized to the threshold value of housekeeping gene GAPDH.

Chromatin immunoprecipitated fragments were amplified by AmpliTaq Gold 360 DNA Polymerase (ThermoFisher) using PTC-1148 MJ Mini Thermal Cycler (Bio-Rad).

The primers used for amplification are indicated in Tables S3 and S4.

### Statistical analyses

Statistical differences were analyzed by Student’s t-test and ANOVA using GraphPad Prism 6.0. When differences were determined to be significant, pairwise comparisons were made by the Tukey test in cases of normal data distribution or Dunn’s test in the opposite case. A 95% confidence level was considered significant.

## Results

### Li^+^ effects on ESC proliferation and GSK3β phosphorylation are dose-dependent

Concentrations of Li^+^ above 2 mM have shown both cytotoxic and anti-proliferative effects in other cell types^16^. ESC were cultured with increasing concentrations of Li^+^ for 3 days to determine the cytotoxicity of Li^+^ on mouse PSCs. 20 mM Li^+^ was toxic for ESCs after 1 day. The number of ESC colonies was considerably fewer than in the other conditions (Figure S1). ESCs treated with 5 and 10 mM Li^+^ showed both lower cell density (Figure S1) and DNA concentration than 2 mM Li^+^ and cells grown in BM (Figure 1a) after 1 day of culture, indicating that concentrations of Li^+^ higher than 2 mM reduce cell proliferation without affecting cell viability.

**Figure 1.**
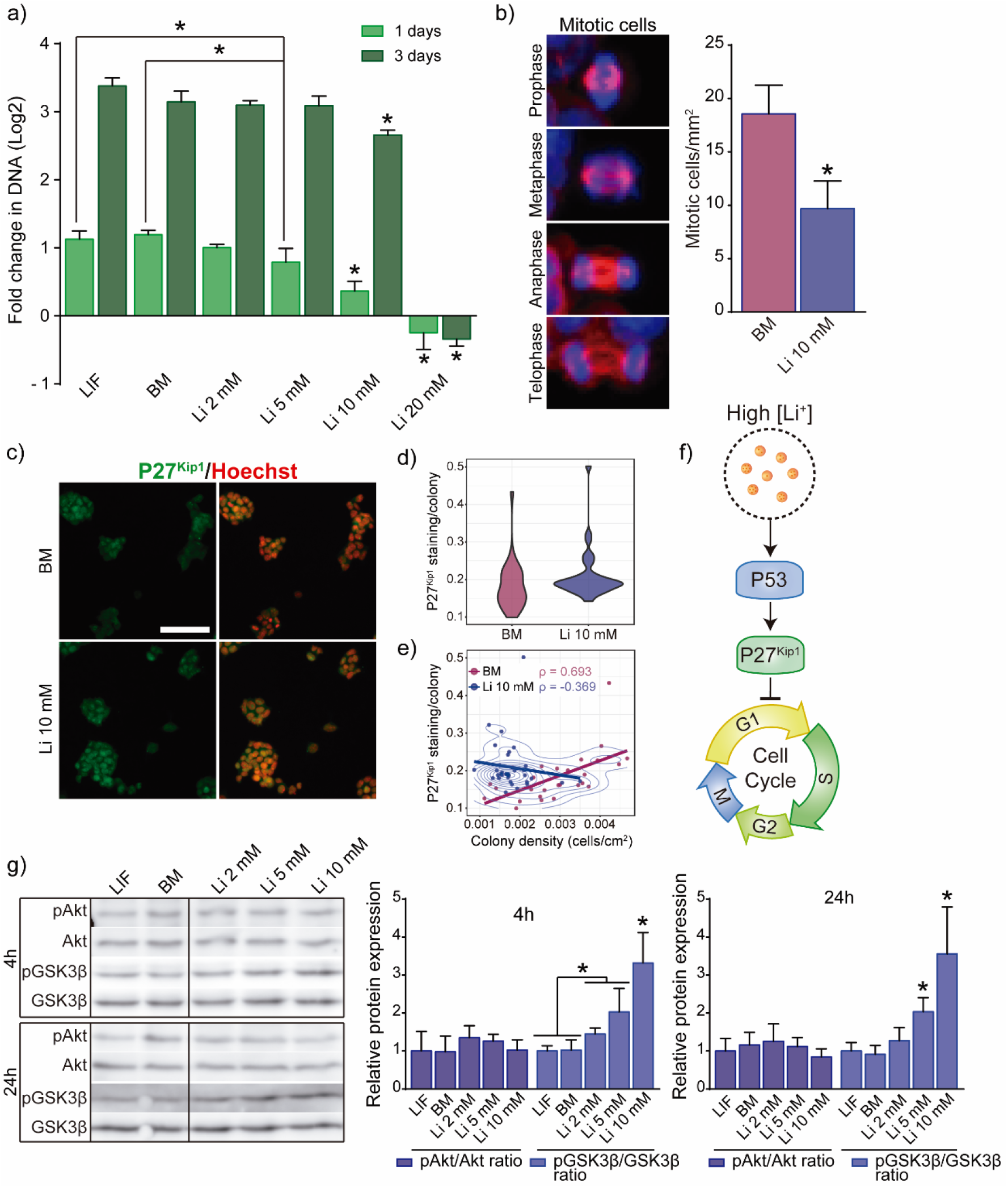
Role of Li^+^ on ESC proliferation and GSK3β phosphorylation. a) DNA concentration in ESCs treated with increasing concentrations of Li^+^ after 1 and 3 days (n = 4). b) Analysis of mitotic cells in ESCs cultured in BM and medium supplemented with 10 mM Li^+^ after 24 h (n = 5). c) Immunofluorescence detection of P27^Kip7^ in ESCs cultured for 24 h in BM and medium supplemented with 10 mM Li^+^. d) Quantification of the nuclear accumulation of P27^Kip1^ per ESCs colony after 24 h. e) Correlation between the nuclear accumulation of P27^Kip1^ in ESC colonies and density of the colonies. Spearman correlation p value was determined. f) Representation of the mechanisms activated by 10 mM Li^+^ to inhibit cell cycle progression. g) Western blot analysis of GSK3β (S9) and Akt (S473) phosphorylation in ESCs treated with increasing concentrations of Li^+^ after 4 and 24 h of culture. GAPDH was used as loading control (n = 4). Graphs show mean ± SD (*p < 0.05).

In order to clarify the mechanism by which Li^+^ reduces ESC proliferation, we studied P53-mediated regulation of cell cycle via P27^Kip1 17^. It has previously been found that 10 mM Li^+^ activates P53 and reduces cell proliferation^18^. We analyzed the number of mitotic cells and the expression of P27^Kip1^ in ESCs cultured in BM and medium supplemented with 10 mM Li^+^. The number of mitotic cells was reduced in the presence of Li^+^ after 24 h (Figure 1b), which agrees with the DNA concentrations shown in Figure 1a. Regarding the P27^Kip1^ expression, ESCs treated with 10 mM Li^+^ showed higher nuclear staining regardless of colony density (Figure 1c, 1d,1f), whereas those cultured in BM showed a monotonical increase of P27^Kip1^ according to colony density (Figure 1e and S2). These results show that the presence of Li^+^ promotes P27^Kip1^expression and arrests the ESC cell cycle.

We next evaluated the Li^+^-mediated phosphorylation of two key PSC regulators: GSK3β and its post-translational regulator Akt^7^ (Figure 1g). Results indicated that GSK3β phosphorylation did not undergo significant changes after 4 and 24 h in cells cultured with LIF and BM. However, for both time points, ESCs supplemented with Li^+^ showed higher dose-dependent GSK3β phosphorylation. For Akt phosphorylation, no significant differences were observed in Li^+^-treated cells, showing similar pAkt/Akt ratio values to those of BM and LIF-cultured cells, suggesting that Li^+^ has no significant effect on Akt phosphorylation (Figure 1g).

### Li^+^ enhances β-catenin expression and nuclear translocation directing ESC differentiation into mesoderm lineage

Although β-catenin activity has been shown to be essential for maintaining ESC integrity^19^, excessive β-catenin accumulation in the nucleus leads to the expression of *Cdx1, Cdx2* and *Brachyury/T* genes implicated in lineage differentiation^20^. To study how Li^+^ affects β-catenin expression and its subcellular distribution, we assessed the nuclear accumulation of β-catenin and GSK3β phosphorylation by immunofluorescence and western blot (Figure 2).

**Figure 2.**
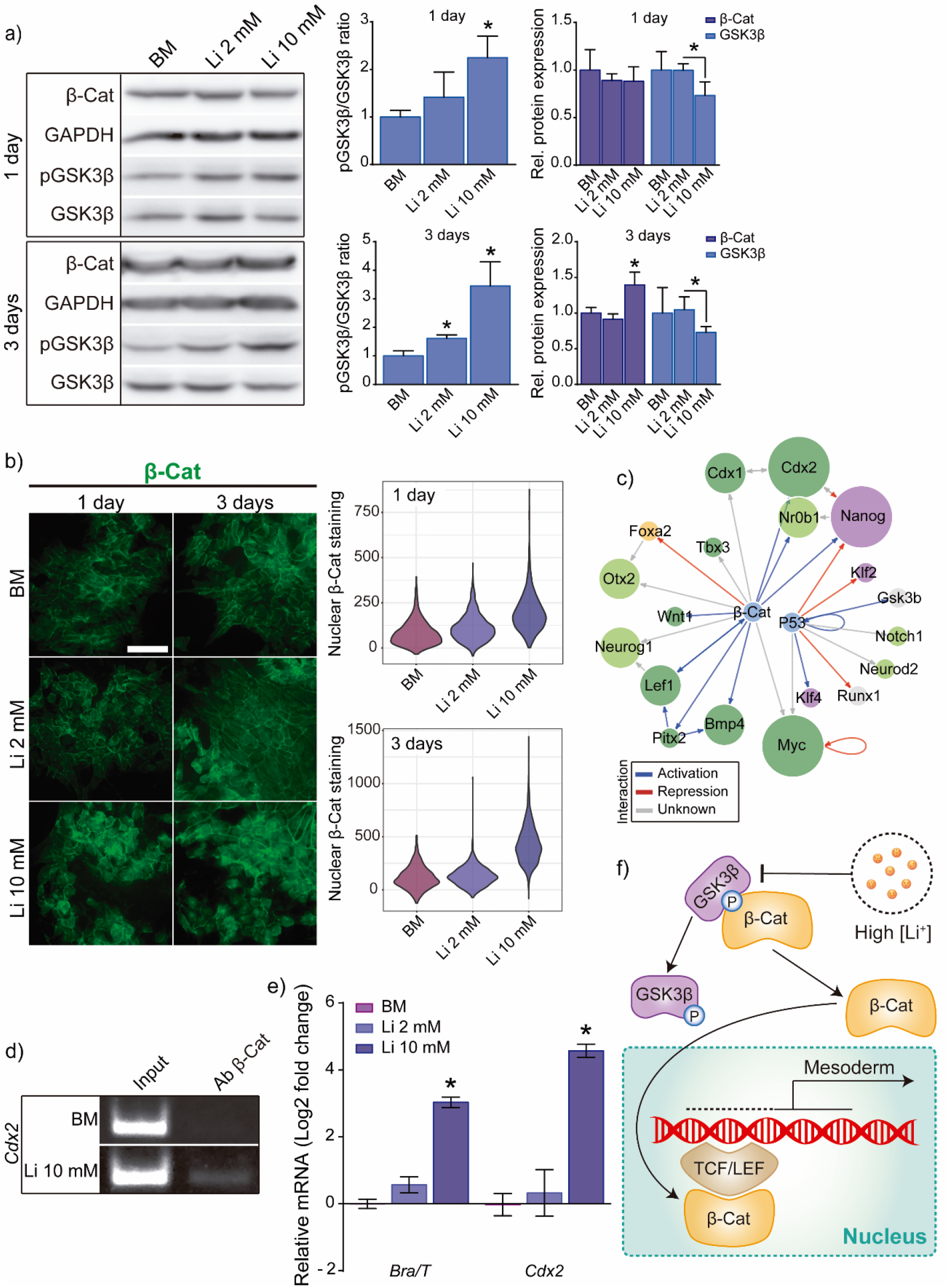
GSK3β activity, β-catenin expression and subcellular localization in Li^+^-treated cells. a) Western blot Analysis of β-catenin expression and pGSK3β/GSK3β ratio in ESCs cultured for 1 and 3 days with different Li^+^ concentrations. GAPDH was used as loading control protein (n = 4). b) Quantification of β-catenin immunostaining after 1 and 3 days of culture. (n > 100 random nuclei (n = 5). c) *In silico* analysis of transcriptional targets of P53 (Tpr53) and β-catenin (Ctnnb1) related to PSC differentiation and self-renewal. Data was obtained from TRRUST data base (V2). d) ChIP on PCR assay of β-catenin binding region in Cdx2 gene. e) qPCR evaluation of mesoderm markers Brachyury/T and Cdx2 expression in ESCs cultured with different concentrations of Li^+^ after 3 days of culture. GAPDH was used as housekeeping gene (n = 4). f) Proposed chain of events in ESCs after exposure to high concentration of Li^+^. Li^+^ triggers GSK3β inactivation and releases β-catenin from the axin complex. Free β-catenin translocates into the nucleus and co-activates the expression of mesoderm differentiation regulators. Graphs show mean ± SD (*p < 0.05).

As expected, after 1 day of culture GSK3β phosphorylation was significantly higher in ESCs treated with 10 mM Li^+^ (as shown in Figure 1g) with no significant differences in terms of β-catenin expression (Figure 2a). After 3 days, the pGSK3β/GSK3β ratio increased monotonically in Li^+^-treated cells, with reduced non-phosphorylated GSK3β expression, whilst β-catenin expression increased. Immunofluorescence analyses showed increased accumulation of β-catenin in the nuclei of ESCs treated with 10 mM Li^+^ after 3 days (Figure 2b). To confirm whether nuclear localization of β-catenin was related to the activation of its transcriptional targets involved in PSC differentiation (Figure 2c and S8), we analyzed β-catenin binding with the *Cdx2* promoter by ChIP on PCR. Only ESCs treated with 10 mM Li^+^ showed amplification in this promoter region (Figure 2d).

To further demonstrate β-catenin transcriptional activity, we analyzed the differential expression of two of its target genes: *Brachyury/T* and *Cdx2*. Both transcription factors are key regulators of the early mesoderm commitment in PSCs^20^. qPCR amplification showed a significantly higher expression of both *Brachyury/T* and *Cdx2* in ESCs treated with 10 mM Li^+^ (Figure 2e). Results indicate that high concentrations of Li^+^ can induce ESC differentiation towards the mesoderm lineage and also that 10 mM Li^+^ can efficiently inhibit the β-catenin destruction complex, following the scheme of events represented in Figure 2f.

### High concentrations of Li^+^ promote ESC differentiation into HE-like cells

Previous studies have shown that GSK3β inhibition induces HE development from ESCs^5,8^. We next studied whether continued exposure to 2 and 10 mM Li^+^ drives ESC differentiation into HE after 6 days of culture. We analyzed pluripotency and HE-specific markers by immunofluorescence and qPCR. The ratio of Oct4 and Sox2 (pluripotency markers) positive cells indicated that ESCs treated with 10 mM Li^+^ showed accelerated differentiation (Figure S3a). After 6 days of culture, HE markers Sca-1, CD31 and Runx1were only present in ESCs cultured with 10 mM Li^+^ (Figure 3). We found that the resulting population of 10 mM Li^+^-treated cells consisted of a combination of either Sca-1+ cells, CD31+ cells or both (Figure 3a and 3b). We further found that continued treatment for 6 days with 10 mM Li^+^ was necessary to reach a significant population of Sca-1+ and CD31+ cells (Figure 3c). When 10 mM Li^+^ was removed from the culture medium (3d-Li 10 mM), the effect on proliferation disappeared for the next 3 days, with significantly higher cell density than in ESCs treated for all 6 days with 10 mM Li^+^, and thus reversing Li’s effect of arresting cell cycle progression. Regarding the expression of Runx1, we observed a faint staining in the nuclei of 10 mM Li^+^-treated cells after 6 days of culture (Figure 3b).

**Figure 3.**
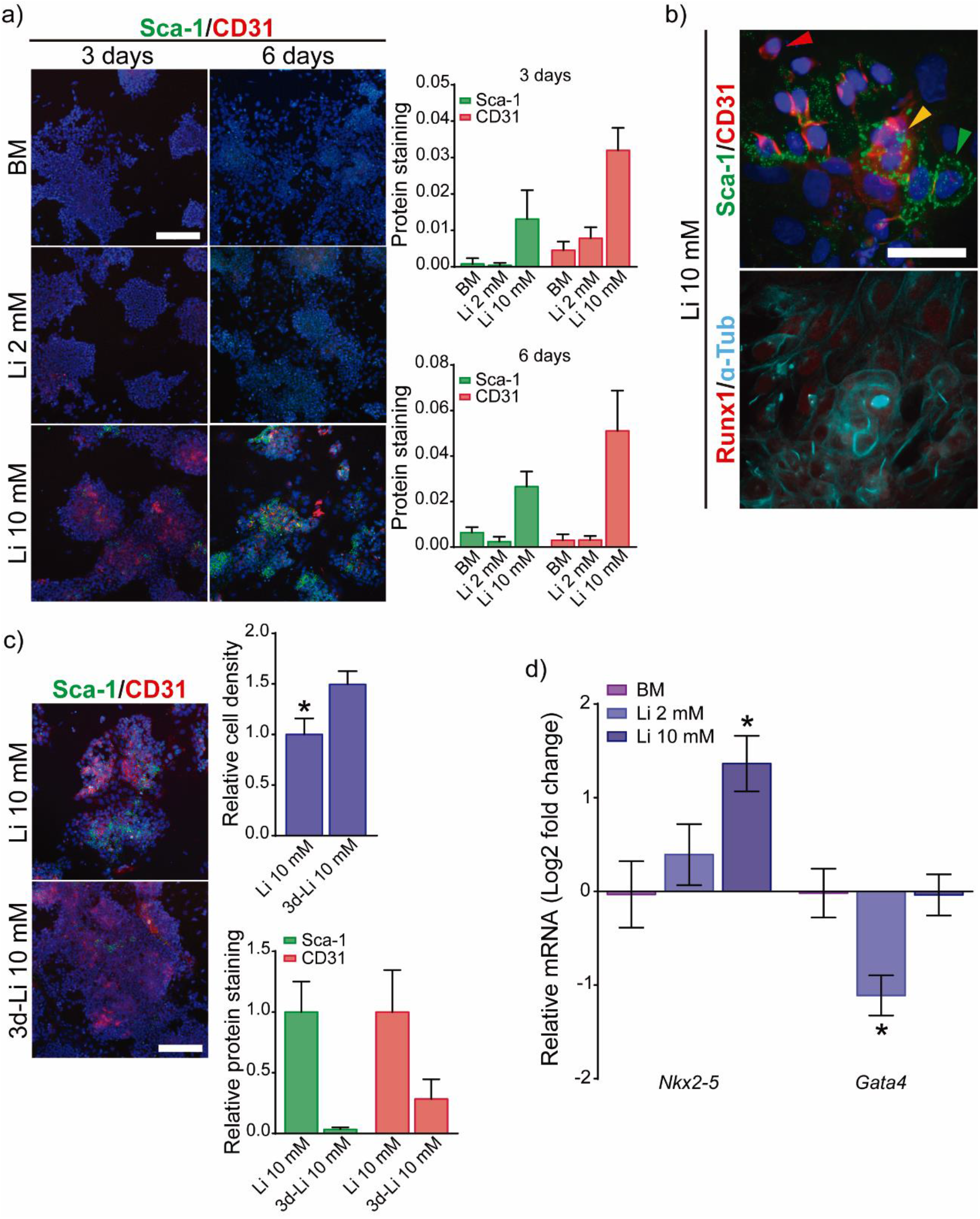
Effect of Li^+^ on ESCs cultured for 6 days in monolayer. a) Immunofluorescence analysis of hemogenic endothelium markers (Sca-1 and CD31) in ESCs cultured for 6 days with different concentrations of Li^+^ (Scale bar: 200 μm) (n = 5). b) Representative images of ESCs cultured with 10 mM Li^+^, with a variety of coexisting CD31^+^ (red arrow), Sca-1^+^ (green arrow), and CD31^+^/Sca-1^+^ (yellow arrow) cells. Immunofluorescence analysis of Runx1 in ESCs treated with 10 mM Li^+^ after 6 days. Scale bar: 50 μm. c) Sca-1 and CD31 expression in ESCs treated for 6 days with 10 mM Li^+^ and further cultured for 3 more days in 10 mM Li^+^ plus 3 days in BM (3 d-10 mM Li^+^). d) qPCR analysis of hemogenic endothelium markers (Nkx2-5 and Gata4) in ESCs cultured with different concentrations of Li^+^ for 6 days (n = 4). Graphs show mean ± SD (*p < 0.05).

We further analyzed the expression of HE precursor markers *Nkx2-5* and *Gata4* after 6 days. *Nkx2-5* was only overexpressed in ESCs treated with 10 mM Li^+^ (Figure 3d). Although no significant differences were found between BM and 10 mM Li^+^-treated ESCs, *Gata4* expression in these conditions was significantly higher than in 2 mM Li^+^-treated cells. *Gata4* expression also characterize ESCs differentiating to early endoderm^21^. To clarify whether this expression is associated with endoderm or mesoderm lineage, we assessed the expression of early endoderm marker Sox17. Only ESCs cultured in BM showed significant Sox17 expression (Figure S3b), indicating that *Gata4* expression of ESCs cultured in BM is related to early endoderm differentiation. Gata4 expression in ESCs treated with 10 mM Li^+^ is therefore due to a mesoderm commitment.

Further analysis of the protein expression of endothelial markers after 6 days of culture revealed an upregulation in β-catenin expression, higher phosphorylation (inactivation) of GSK3β and expression of VE-cadherin in ESCs supplemented with 10 mM Li^+^ (Figure S3c), supporting the hypothesis that 10mM Li^+^ induces HE development from ESCs via GSK3β phosphorylation and β-catenin activation.

### Li^+^-induced ESCs-derived Hemogenic Endothelium (HE) cells can be maturated to obtain Hematopoietic stem cells (HSCs)

HSCs deriving from HE arise from a subset of population characterized by the expression of both Runx1+ and Sca-1+ cells^22,23^. We observed that after 6 days of culture with 10 mM Li^+^, ESCs successfully differentiated into Sca-1+ cells and showed a faint expression of Runx1 and nuclear β-catenin (Figure S4). To further investigate the potential of Sca-1+ cells derived from ESCs to differentiate into HSCs, we induced these cells to differentiate under maturation medium N2B27 for 5 and 11 days (Figure 4a). The expression of HE (Sca-1, CD31, Runx1 and Sox17) and HSCs (Sox17, Runx1, CD34 and CD166) markers were analyzed by immunostaining after 5 and 11 days of maturation in N2B27 maturation medium. After 5 days, HE markers Runx1 and Sox17 were upregulated for the ESCs pre-treated with 10 mM Li^+^, observing an increasing number of cells co-expressing both markers (Figure 4b and 4c). These cells begin to appear around Sca-1-expressing colonies (Figure 4b and S5b), suggesting their origin lies in Sca-1+ cells, which reduce Sca-1+ levels after maturation and increase Rux1 and Sox17 markers, indicating HSC formation.

**Figure 4.**
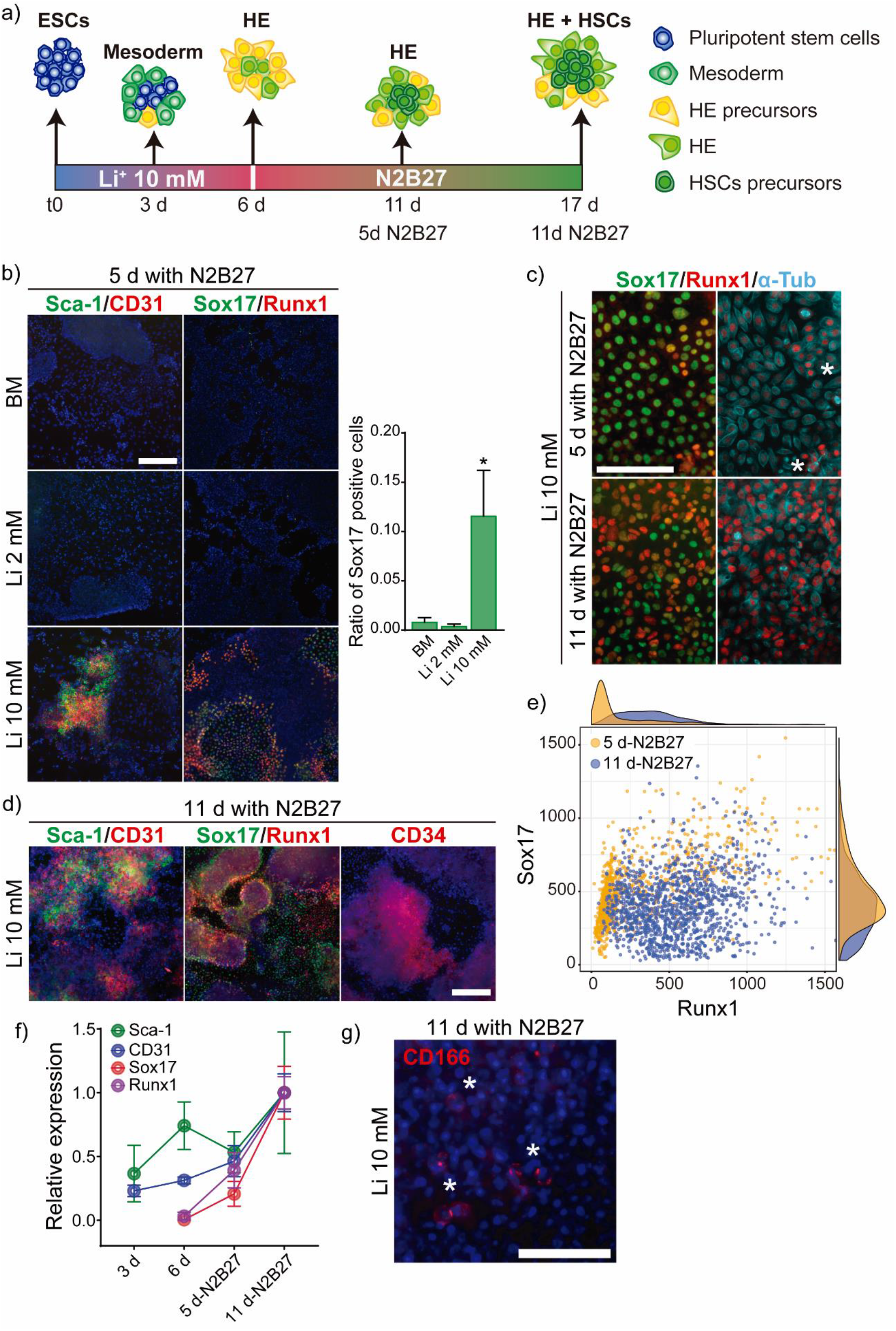
Maturation of HE cells derived from ESCs treated with 10 mM Li^+^. a) Experimental set up for differentiation of ESCs in presence of 10 mM Li^+^ for 6 days and then maturated in HSC precursors for 11 days under differentiation conditions (maturation medium N2B27). b) Immunofluorescence detection of HE (Sca-1 and CD31) and HSCs (Sox17 and Runx1) markers in ESCs pre-cultured for 6 days in presence of different concentrations of Li^+^ and then maturated for 5 days in N2B27 (n = 5). Scale bar: 200 μm. c) Immunofluorescence images of the evolution of Sox17 and Runx1 expression during the maturation of ESCs pre-cultured with 10 mM Li^+^. Cell clamping indicated by white asterisks. Scale bar: 200 μm. d) Immunofluorescence detection of HE (Sca-1 and CD31) and foetal (Sox17) and adult (CD34, Runx1) HSCs markers in ESCs pre-cultured for 6 days with 10 mM Li^+^ and then maturated for 11 days in N2B27. Scale bar: 200 μm. e) Analysis of the nuclear accumulation of RunX1 and Sox17 during the maturation of ESCs pre-cultured for 6 days with 10 mM Li^+^ and then maturated for 5 and 11 days in N2B27 (n > 1000 nuclei). f) Analysis of Sca-1, CD31, Sox17and Runx1 during HE development and maturation into HSC progenitors (n = 5). g) Immunostaining of HSC marker CD166 (white asterisks) in ESCs pre-cultured for 6 days with 10 mM Li^+^ and then maturated for 11 days in N2B27. Scale bars: 50 μm. Graphs show mean ± SD(*p < 0.05).

Bright field images showed that ESCs pre-treated with 10 mM Li^+^ and matured for 7 days possessed both endothelial-like cell and clustered and rounded cell morphologies (Figure S5a), reinforcing the idea of hematopoietic commitment. After 11 days cell morphologies of ESCs pre-cultured in BM or 2 mM Li^+^ and those treated with 10 mM Li^+^ were clearly different. Whilst 10 mM Li^+^-treated ESCs showed a significant population of rounded cells over endothelial-like colonies, the other conditions showed many neural-like cells (Figure S5a). The CD31+, Sca-1+, Runx1+ and Sox17+ cell population increased even more in ESCs pre-treated with 10 mM Li^+^ after 11 days in a maturation medium, with a large number of clamped colonies (Figure 4d and 4f). Indeed, after 11 days the Runx1/Sox17 ratio in the cells that co-express Runx1 and Sox17 was higher than the values obtained 6 days before (Figures 4e, S5b and c). By contrast, the remaining cell population had colonies expressing HSCs marker CD34+ and some CD166+ cells (Figure 4d and 4g). After 11 days of culture under maturation conditions we also observed a gradual change of morphology, forming clusters composed of rounded cells, supporting the idea of endothelial-to-hematopoietic transition (EHT) (Figure S5c).

To further test our hypothesis based on Li^+^‘s capacity to induce ESC differentiation into HE and further maturation into HSCs we analyzed relevant parameters after culturing ESC in suspension (Figure S6 and S7), obtaining spheroids and showing that Li^+^ successfully induces ESC differentiation into HE.

## Discussion

GSK3β activity is an essential molecular cue controlling ESCs’ fate^20^,^24^. Previous reports have indicated that GSK3 activity is related to both ESC self-renewal/differentiation^20^ and β-catenin inhibition^7^. Other works related to Li^+^ effects suggest that high concentrations of Li^+^ inhibited ESC differentiation towards cardiac lineage, favoring neural differentiation^10^. Despite these reports, we did not find any neural (Figure S7c) or cardiac (Figure S8) differentiation when using 10 mM Li^+^. In fact, we have found that the presence of Li^+^ did not affect ESCs’ pluripotency potential (Figures S3a), and that the continued exposure of ESCs to 10 mM Li^+^ strongly inhibited (phosphorylated) GSK3β and induced β-catenin transcriptional activation (Figure 2), giving rise to mesoderm development (Figure 2) and subsequent HE specification (Figure 3), in ESCs cultured in both monolayer (Figure 3 and 4) and in suspension (Figures S6 and S7), regardless of the composition of the culture medium.

Our findings are supported by other studies based on the use of GSK3β inhibitors^8^. The activation of Wnt signaling by either Wnt or GSK3β inhibitors has shown to induce HE differentiation in ESCs^5,8,9^ Here, we demonstrate that after 6 days of culture, initial ESC colonies expressed Sca-1, CD31, VE-cadherin, Nkx2-5 and a Runx1 (Figure 3 and S3). These markers are characteristic of an endothelium with the potential to develop HSCs^2,25^ Furthermore, *in silico* analysis predicts that the combined activity of both P53 and β-catenin activated by 10 mM Li^+^ could target many genes related to mesodermal lineages differentiation (Figure 2c) and HE development (Figure S9). Even though Nkx2-5 is a master regulator of cardiogenesis^26^, its expression has also been found in hemogenic endothelial cells^25^. *In silico* analysis also predicted that the P53 activation by 10 mM Li^+ 18^ could hinder the development of mature HE cells because of the transcriptional inactivation of Runx1 (Figure 2c), which is essential for the development of HSC precursors from HE cells^27^,^28^. Should this hypothesis be valid, it would explain the faint staining of Runx1 observed after 6 days in the presence of 10 mM Li^+^ and the rapid increase when the medium was replaced by the maturation medium (without Li^+^).

The hemogenic capability of endothelial cells obtained with 10 mM of Li^+^ was confirmed after their defined maturation (Figure 4). After 11 days in N2B27 maturation medium, we found rounded cells expressing Runx1 and Sox17. As Sox17 expression is a feature of fetal HSCs before the acquisition of the adult phenotype^29^, HSCs developed from PSCs can be expected to share many characteristics with fetal HSCs because of their embryonic-like origin^30^. Furthermore, at 11 days in maturation medium, the ratio between Runx1/Sox17 was significantly higher (Figure 4c and 4e) than the values observed at 5 days. As Sox17 has been reported to impair Runx1 transcriptional activity, its downregulation is necessary for EHT^27^. These results were reinforced by the changes in the cell morphologies; we observed Runx1/Sox17 expression in clumped and rounded cells (Figure S5c), while Runx1 expression was significantly higher than that of Sox17 (Figure 4c), indicating EHT. HSC development was suggested by the multiplication of CD34+ colonies and CD166-expressing cells (Figure 4d and 4g)^31^.

Altogether, we have defined the mechanisms activated by lithium on murine ESC self-renewal and differentiation. Li^+^ can thus be regarded as a powerful molecule capable of simply and efficiently activating the P53 and β-catenin transcription factors involved in HE development from ESC. We have also shown the ability of these lithium-treated ESCs to further derive into HSCs after defined maturation, confirming the endothelial to hematopoietic transition. These results represent a novel strategy for successfully generating HSC *in vitro* as a multipotent source of stem cells for biomedical blood and muscle applications.

## Acknowledgements

Funding: PR acknowledges support from the Spanish Ministry of Science, Innovation and Universities (RTI2018-096794), and Fondo Europeo de Desarrollo Regional (FEDER). CIBER-BBN is an initiative funded by the VI National R&D&I Plan 2008-2011, Iniciativa Ingenio 2010, Consolider Program, CIBER Actions and financed by the Instituto de Salud Carlos III with assistance from the European Regional Development Fund.

MSS acknowledges support from the UK Engineering and Physical Sciences Research Council (EPSRC - EP/P001114/1).

## References

1. Ng AP, Alexander WS. Haematopoietic stem cells: past, present and future. Cell Death Discov. 2017;3(1):17002.

2. Ackermann M, Liebhaber S, Klusmann J, Lachmann N. Lost in translation: pluripotent stem cell-derived hematopoiesis. EMBO Mol. Med. 2015;7(11):1388–1402.

3. Hirschi KK. Hemogenic endothelium during development and beyond. Blood. 2012;119(21):4823–4827.

4. Swiers G, Rode C, Azzoni E, De Bruijn MFTR. A short history of hemogenic endothelium. Blood Cells, Mol. Dis. 2013;51(4):206–212.

5. Sturgeon CM, Ditadi A, Awong G, Kennedy M, Keller G. Wnt signaling controls the specification of definitive and primitive hematopoiesis from human pluripotent stem cells. Nat. Biotechnol. 2014;32(6):554–561.

6. Lian X, Hsiao C, Wilson G, et al. Robust cardiomyocyte differentiation from human pluripotent stem cells via temporal modulation of canonical Wnt signaling. Proc. Natl. Acad. Sci. 2012;109(27):E1848–E1857.

7. Wu D, Pan W. GSK3: a multifaceted kinase in Wnt signaling. Trends Biochem. Sci. 2010;35(3):161–168.

8. D’Souza SS, Maufort J, Kumar A, et al. GSK3β Inhibition Promotes Efficient Myeloid and Lymphoid Hematopoiesis from Non-human Primate-Induced Pluripotent Stem Cells. Stem Cell Reports. 2016;6(2):243–256.

9. Freland L, Beaulieu J-M. Inhibition of GSK3 by lithium, from single molecules to signaling networks. Front. Mol. Neurosci. 2012;5(February):1–7.

10. Schmidt MM, Guan K, Wobus AM. Lithium influences differentiation and tissue-specific gene expression of mouse embryonic stem (ES) cells in vitro. Int. J. Dev. Biol. 2001;45(2):421–429.

11. Chen T, Wang F, Wu M, Wang ZZ. Development of Hematopoietic Stem and Progenitor Cells From Human Pluripotent Stem Cells. J. Cell. Biochem. 2015;116(7):1179–1189.

12. Esiashvili N, Pulsipher MA. Hematopoietic stem cell transplantation. Pediatr. Oncol. 2018;(9783319435442):301–311.

13. Ruiz-Herguido C, Guiu J, D’Altri T, et al. Hematopoietic stem cell development requires transient Wnt/β-catenin activity. J. Exp. Med. 2012;209(8):1457–1468.

14. Strahl-Bolsinger S, Hecht A, Luo K, Grunstein M. SIR2 and SIR4 interactions differ in core and extended telomeric heterochromatin in yeast. Genes Dev. 1997;11(1):83–93.

15. Pfaffl MW. A new mathematical model for relative quantification in real-time RT-PCR. Nucleic Acids Res. 2001;29(9):e45.

16. Allagui MS, Vincent C, El feki A, Gaubin Y, Croute F. Lithium toxicity and expression of stress-related genes or proteins in A549 cells. Biochim. Biophys. Acta - Mol. Cell Res. 2007;1773(7):1107–1115.

17. Zhang HM, Yuan J, Cheung P, et al. Gamma Interferon-Inducible Protein 10 Induces HeLa Cell Apoptosis through a p53-Dependent Pathway Initiated by Suppression of Human Papillomavirus Type 18 E6 and E7 Expression. Mol. Cell. Biol. 2005;25(14):6247–6258.

18. Mao CD, Hoang P, DiCorleto PE. Lithium Inhibits Cell Cycle Progression and Induces Stabilization of p53 in Bovine Aortic Endothelial Cells. J. Biol. Chem. 2001;276(28):26180–26188.

19. Raggioli A, Junghans D, Rudloff S, Kemler R. Beta-catenin is vital for the integrity of mouse embryonic stem cells. PLoS One. 2014;9(1):e86691.

20. Chen Y, Blair K, Smith A. Robust self-renewal of rat embryonic stem cells requires fine-tuning of glycogen synthase kinase-3 inhibition. Stem Cell Reports. 2013;1(3):209–217.

21. Keller G. Embryonic stem cell differentiation: emergence of a new era in biology and medicine. Genes Dev. 2005;19(10):1129–55.

22. Valenta T, Hausmann G, Basler K. The many faces and functions of β-catenin. EMBO J. 2012;31(12):2714–2736.

23. Yzaguirre AD, Speck NA. Insights into blood cell formation from hemogenic endothelium in lesser-known anatomic sites. Dev. Dyn. 2016;245(10):1011–1028.

24. Martello G, Sugimoto T, Diamanti E, et al. Esrrb is a pivotal target of the Gsk3/Tcf3 axis regulating embryonic stem cell self-renewal. Cell Stem Cell. 2012;11(4):491–504.

25. Zamir L, Singh R, Nathan E, et al. Nkx2.5 marks angioblasts that contribute to hemogenic endothelium of the endocardium and dorsal aorta. Elife. 2017;6:1–31.

26. Akazawa H, Komuro I. Cardiac transcription factor Csx/Nkx2-5: Its role in cardiac development and diseases. Pharmacol. Ther. 2005;107(2):252–268.

27. Lizama CO, Hawkins JS, Schmitt CE, et al. Repression of arterial genes in hemogenic endothelium is sufficient for haematopoietic fate acquisition. Nat. Commun. 2015;6(May):1–10.

28. Chen MJ, Yokomizo T, Zeigler BM, Dzierzak E, Speck NA. Runx1 is required for the endothelial to haematopoietic cell transition but not thereafter. Nature. 2009;457(7231):887–891.

29. Kim I, Saunders TL, Morrison SJ. Sox17 Dependence Distinguishes the Transcriptional Regulation of Fetal from Adult Hematopoietic Stem Cells. Cell. 2007;130(3):470–483.

30. Hanif MA, Bhatti HN, Ali MA, Asgher M, Bhatti IA. Heavy metals tolerance and biosorption potential of white rot fungi. Asian J. Chem. 2010;22(1):335–345.

31. Chitteti BR, Poteat BA, Cardoso AA, et al. CD166 regulates human and murine hematopoietic stem cells and the hematopoietic niche. Blood. 2014;124(4):519–529.

