## supplementary information for "Lithium directs embryonic stem cell differentiation versus hemogenic endothelium"

#### 1. Supplementary materials and methods

#### Serum free differentiation of ESCs

ESCs were cultured in suspension for 2 days in a serum-free (SF: DMEM/F12 with N2 and B27 supplements) medium for mesoderm differentiation. After incubation, spheroids were collected and the SF medium was supplemented with differentiation factors: 10 µM Activin A, 10 µM BMP4 and 10 µM VEGF (Peprotech). After 4 days of culture under differentiation conditions, spheroids were disaggregated using 0.025% Trypsin/EDTA (Gibco) and further seeded on fibronectin coated plates (10 µg/mL, Sigma-Aldrich) to allow differentiated cells to spread for subsequent analysis. Cells were cultured in SF medium supplemented with differentiation factors for 2 additional days.

#### Histological sections

For fluorescence immunodetection EBs were fixed with 4% formaldehyde for 30 min at room temperature. Then, EBs were embedded into low temperature gelling agarose (Sigma-Aldrich) and included in OCT (Tissue-Tek) for 15 µm cryosections obtained on a cryostat CM1520 (Leica). OCT sections were immunostained as previously described.

For histological stainings, EBs were fixed with 4% formaldehyde overnight at 4° C. EBs were embedded into low gelling-temperature agarose (Sigma-Aldrich) and paraffin (Polyester Wax, Electron Microscopy) for 6 µm histological sections obtained with microtome HM 350 S (ThermoFisher). Paraffin sections were stained with Hematoxylin/Eosin (Sigma-Aldrich), as previously described (Fischer et al., 2008).

**Table 1.** Antibodies for western blot.

| Primary antibodies | Dilution | Reference |
|---|---|---|
| Goat anti-Oct4 | 1/400 | SantaCuz Biotechnology |
| Rabbit anti-Sox2 | 1/500 | ThermoFisher |
| Rabbit anti-P27$^{Kip1}$ | 1/200 | ThermoFisher |
| Goat anti-Sox17 | 1/500 | R&D systems |
| Rat anti-Sca1 (clone: E13 161-7) | 1/1000 | Abcam |
| Rbbit anti-CD31 | 1/300 | Abcam |
| Rabbit anti-CD34 | 1/300 | Abcam |
| Rabbit anti-αSMA | 1/500 | Abcam |
| Rabbit anti-Runx1 | 1/500 | Abcam |
| Mouse anti-Sarcomeric α-actinin | 1/400 | Abcam |
| Rabbit anti-β Catenin | 1/500 | ThermoFisher |
| **Secondary antibodies** | **Dilution** | **Reference** |
| Alexa 488 conjugated goat anti-mouse | 1/700 | ThermoFisher |
| Alexa 555 conjugated goat anti-mouse | 1/700 | ThermoFisher |
| Alexa 488 conjugated goat anti-rabbit | 1/700 | ThermoFisher |
| Alexa 555 conjugated goat anti-rabbit | 1/700 | ThermoFisher |
| Alexa 633 conjugated goat anti-rabbit | 1/300 | ThermoFisher |
| DyLight 488 conjugate donkey anti-goat IgG | 1/700 | ThermoFisher |

**Table 2.** Antibodies and dilutions used for western blot experiments.

| Primary antibodies | Dilution | Reference |
|---|---|---|
| Rabbit anti-Akt (Clone J.314.4) | 1/1500 | ThermoFisher |
| Mouse anti-phospho Akt-serine 473 (Clone 6F5) | 1/1500 | Millipore |
| Rabbit anti-VE cadherin | 1/1000 | Abcam |
| Rabbit anti-β Catenin | 1/500 | ThermoFisher |
| Rabbit anti-GSK3β (Clone 3D10) | 1/1500 | ThermoFisher |
| Rabbit anti-phospho GSK3β-serine 9 (Clone C.367.3) | 1/1000 | ThermoFisher |
| Rabbit anti-GAPDH | 1/3000 | ThermoFisher |
| **Secondary antibodies** | **Dilution** | **Reference** |
| Donkey anti-mouse HRP conjugate | 1/10000 | GE Healthcare |
| Donkey anti-rabbit HRP conjugate | 1/10000 | GE Healthcare |

**Table 3.** RT-qPCR primer sequences.

| Primers | | Sequence |
|---|---|---|
| **Brachyury/T** | 5’-Forward | GGTGGCTTGTTCCTGGTGC |
| | 5’-Reverse | GTAGGTGGGCTGGCGTTAT |
| **Cdx2** | 5’-Forward | AAACCTGTGCGAGTGGATG |
| | 5’-Reverse | TCTGTGTACACCACCCGGTA |
| **Gata4** | 5’-Forward | CCTAAACCTTACTGGCCGTAGC |
| | 5’-Reverse | ACAATGTTAACGGGTTGTGGAG |
| **Nkx2-5** | 5’-Forward | CAAGTGCTCTCCTGCTTTCC |
| | 5’-Reverse | GGCTTTGTCCAGCTCCACT |
| **GAPDH** | 5’-Forward | AGGTCGGTGTGAACGGATTTG |
| | 5’-Reverse | TGTAGACCATGTAGTTGAGGTCA |

**Table 4.** ChIP on PCR primer sequences.

| Primers | | Sequence |
|---|---|---|
| **Cdx2** | 5'-Forward | AAACCTGTGCGAGTGGATG |
| | 5'-Reverse | TCTGTGTACACCACCCGGTA |

### 2. Supplementary results

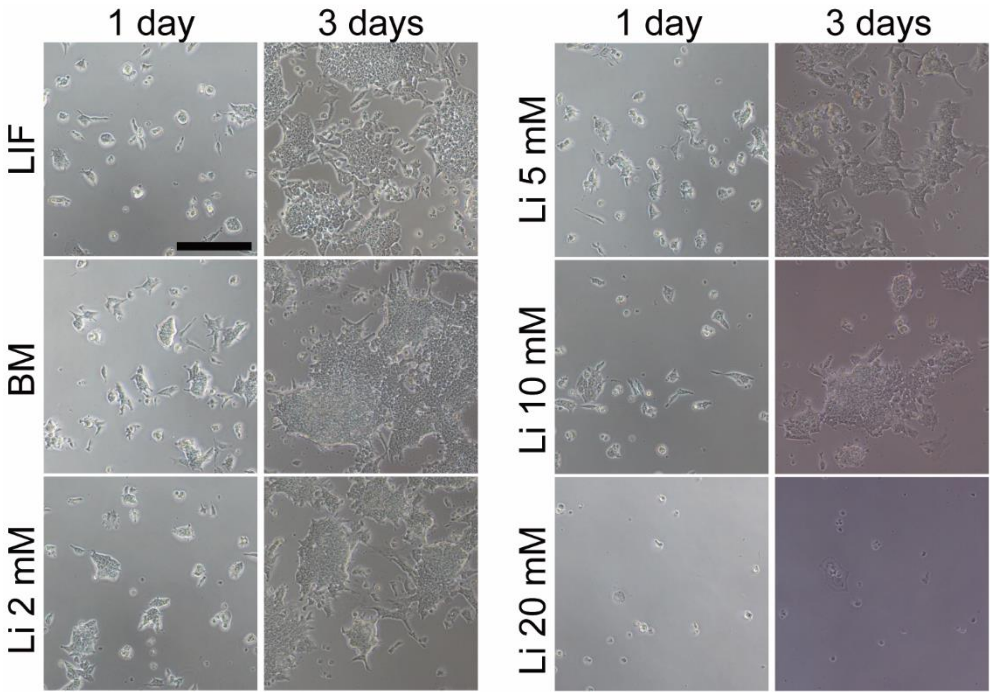


**Supplementary Figure 1**. Role of lithium in ESCs survival. (Extended figure 1a).

Bright field images of ESCs cultured with different concentrations of $Li^{+}$ after 1 and 3 days of culture (Scale bar: 100 µm).

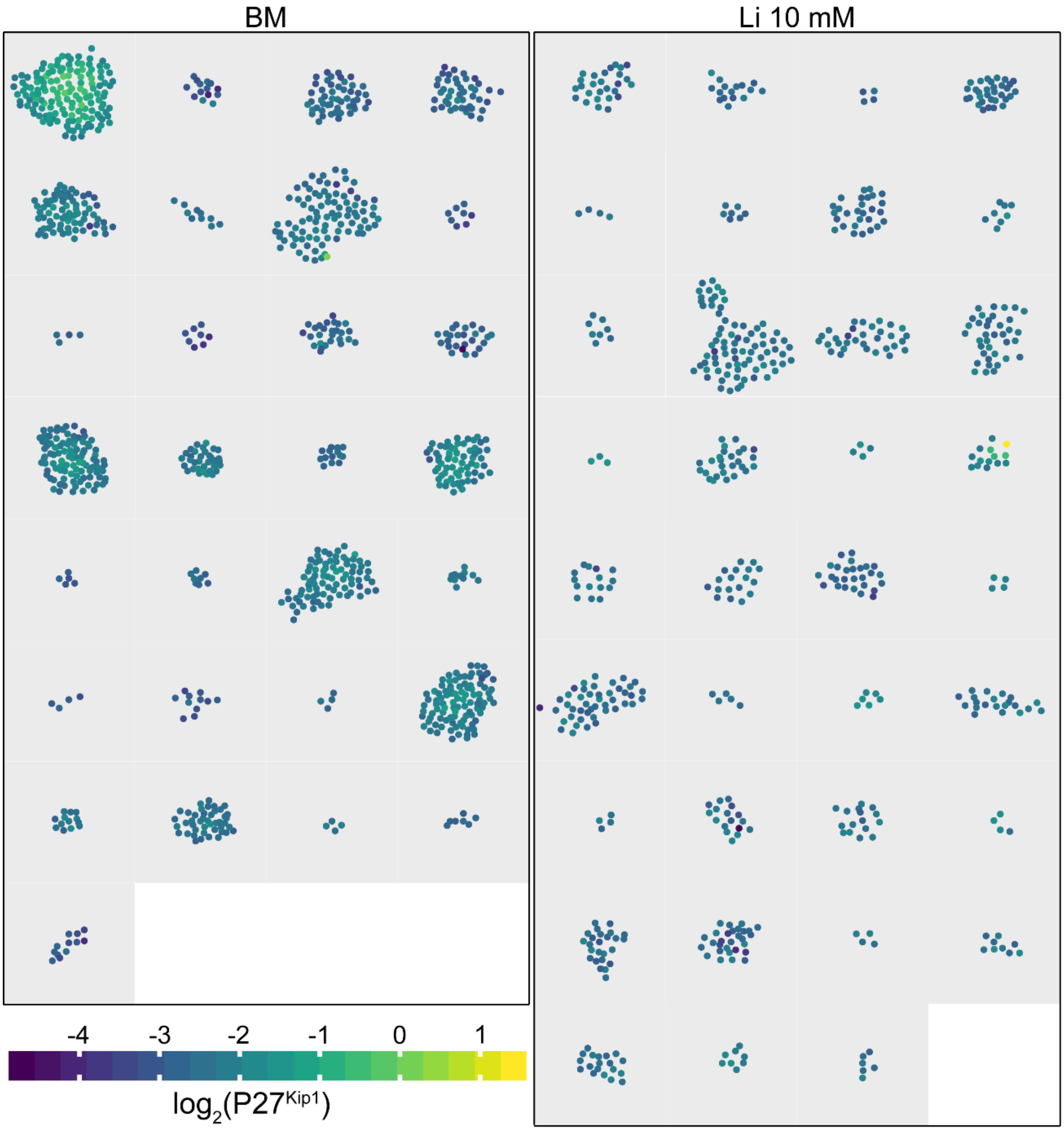


**Supplementary Figure 2**. $Li^+$ reduces ESC proliferation (Extension of Figure 1c).

Representation of several ESC colonies cultured for 24h in BM and BM supplemented with 10 mM $Li^+$. Each nucleus in the colony was depicted individually and stained according to the levels of $P27^{Kip1}$.

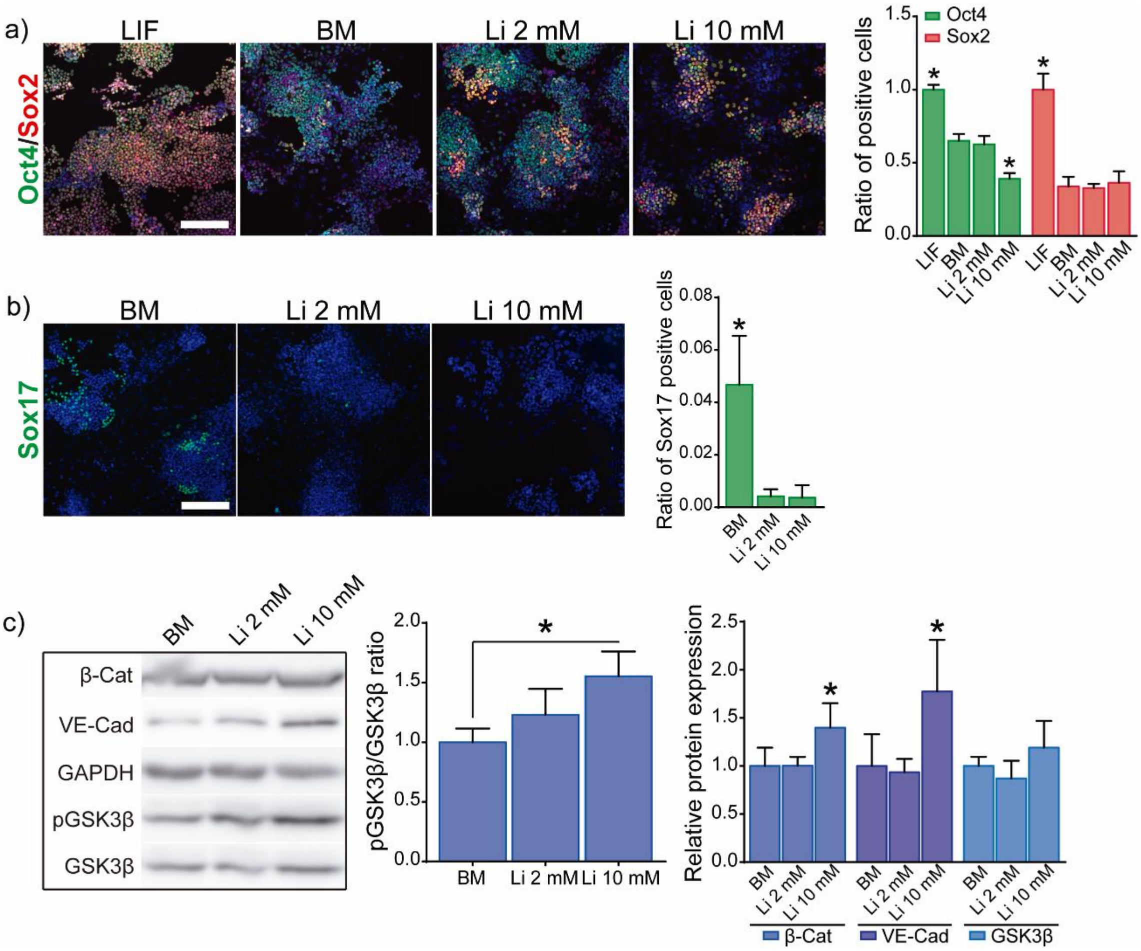


**Supplementary Figure 3**. Role of $Li^+$ in ESC self-renewal and expression of differentiation markers after 6 days and 1 passage (after 3 days, extended Figure 3).

a) Immunofluorescence analysis of Oct4 and Sox2 pluripotency markers in ESCs cultured for 6 days in BM and medium supplemented with different concentrations of $Li^+$ (n = 5). Scale bar: 200 µm.

b) Immunofluorescence analysis of endoderm lineage markers (Sox17) in ESCs cultured for 6 days in BM and medium supplemented with different concentrations of $Li^+$. Scale bar: 200 µm.

c) Western blot analysis of GSK3β (S9) phosphorylation, expression of β-catenin and VE-cadherin in ESCs treated with different concentrations of $Li^+$ after 6 days. GAPDH was used as loading control protein (n = 4).

Graphs show mean ± standard deviation. Significant differences were determined by ANOVA test; *$p < 0.05$.

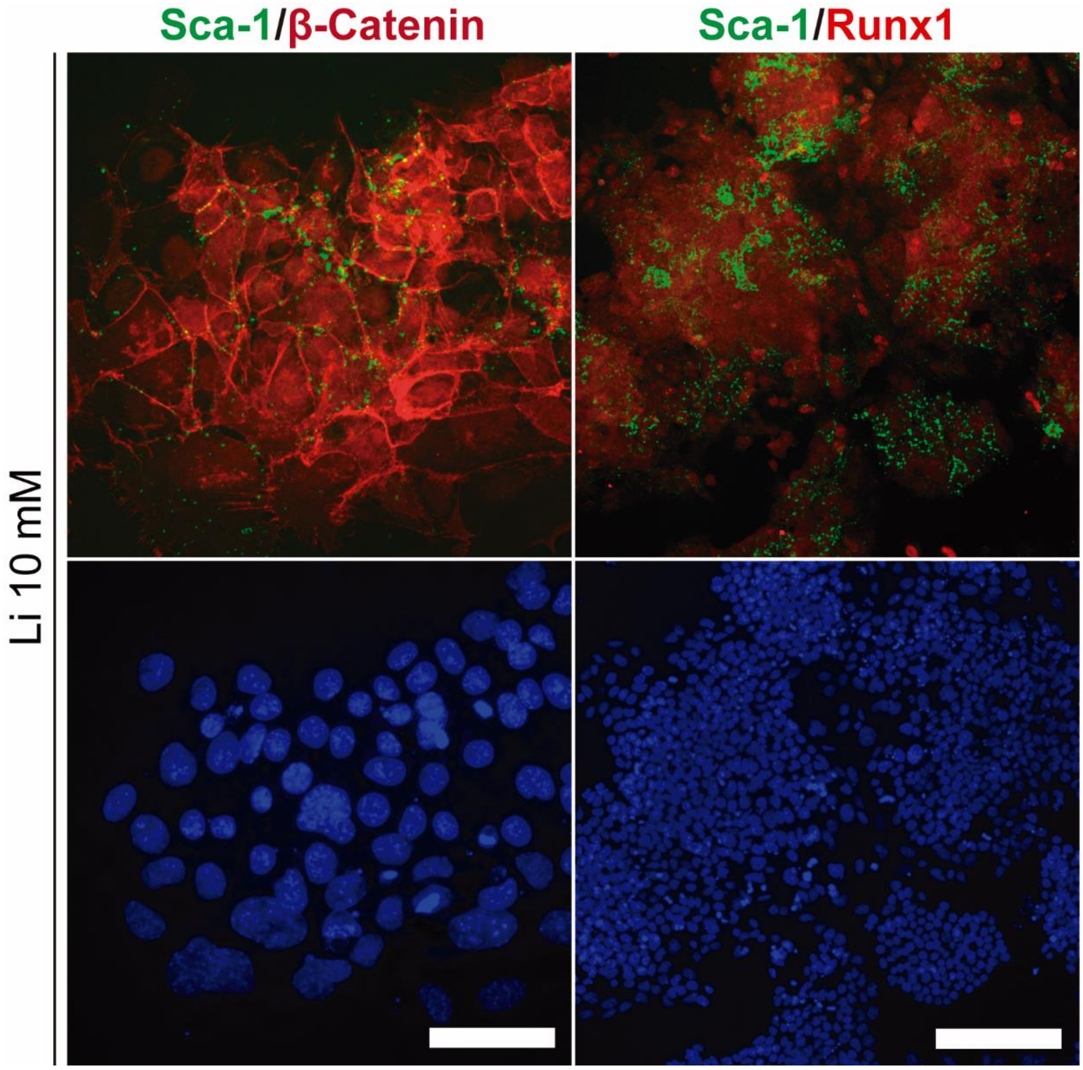


**Supplementary Figure 4**. Role of 10 mM $Li^+$ in the activation of hemogenic endothelial differentiation in ESCs after 6 days. (Extended Figure 3).

Immunofluorescence analysis of the co-expression of Sca-1/β-catenin (scale bar: 50 µm) and Sca-1/Runx1 (scale bar: 200 µm) in ESCs cultured for 6 days in medium supplemented with 10 mM $Li^+$.

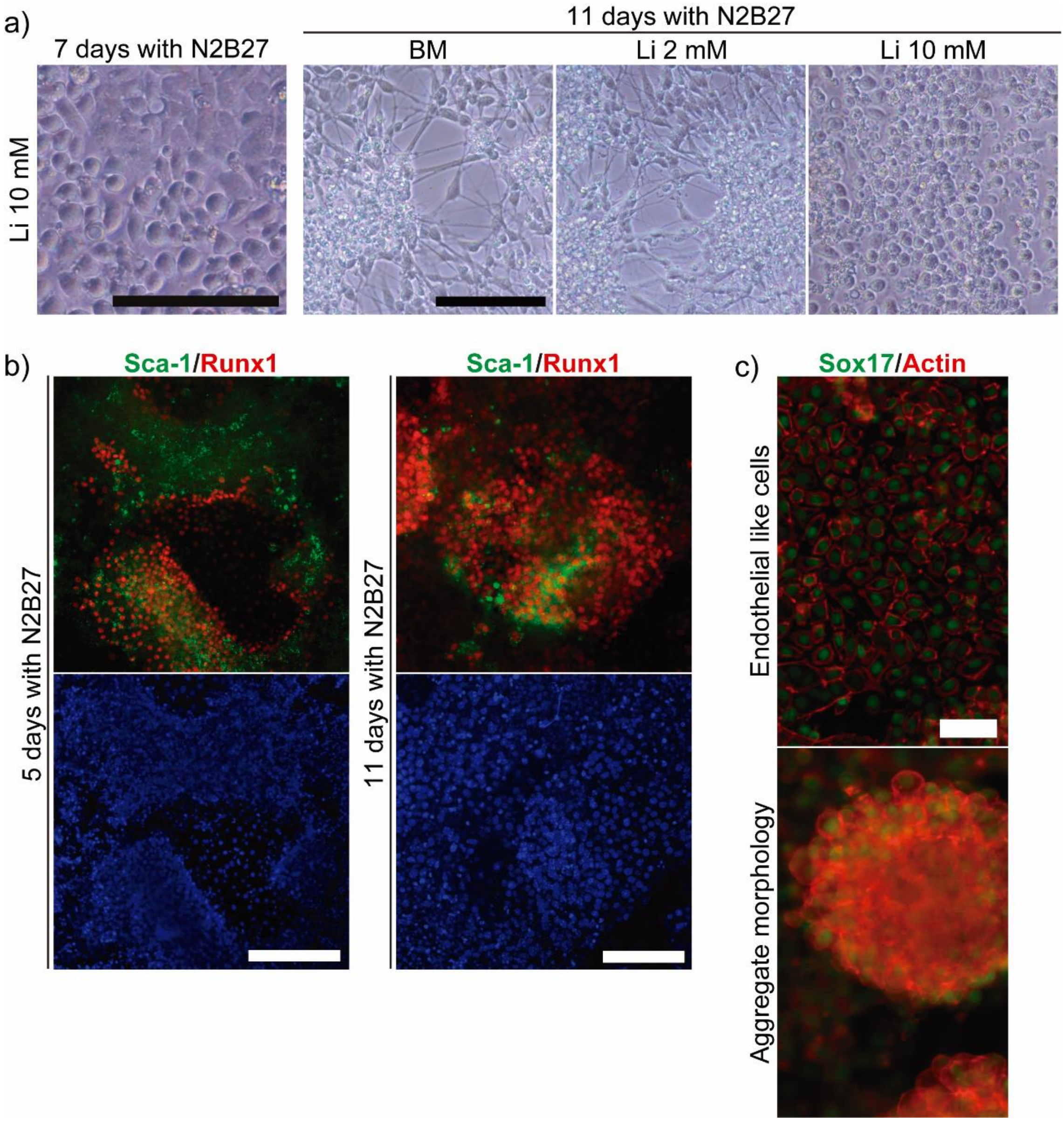


**Supplementary Figure 5**. Maturation of hemogenic endothelial cells derived from ESCs treated with 10 mM Li+. (Extended Figure 4).

a) Bright field images of cell morphology in ESCs pre-cultured in presence of 10 mM $Li^+$ and then maturated with N2B27 for 7 days. Both endothelial cell-like morphology and the formation of rounded cell clusters can be seen (Scale bar: 100 µm).

Bright field images of cell morphology of ESCs pre-treated with different concentrations of $Li^+$ and then maturated for 11 days in N2B27 medium (Scale bar: 100 µm).

b) Immunofluorescence detection of markers related to hemogenic endothelium lineage (Sca-1 and Runx1) in ESCs pre-cultured in presence of 10 mM $Li^{+}$ and then maturated with N2B27 for 5 days (Scale bar: 200 µm) and 11 days (Scale bar: 100 µm).

c) Immunofluorescence analysis of the morphology of the cells expressing Sox17 (hemogenic endothelial and hematopoietic stem cell precursor marker) after 11 days of maturation. Scale bar: 50 µm.

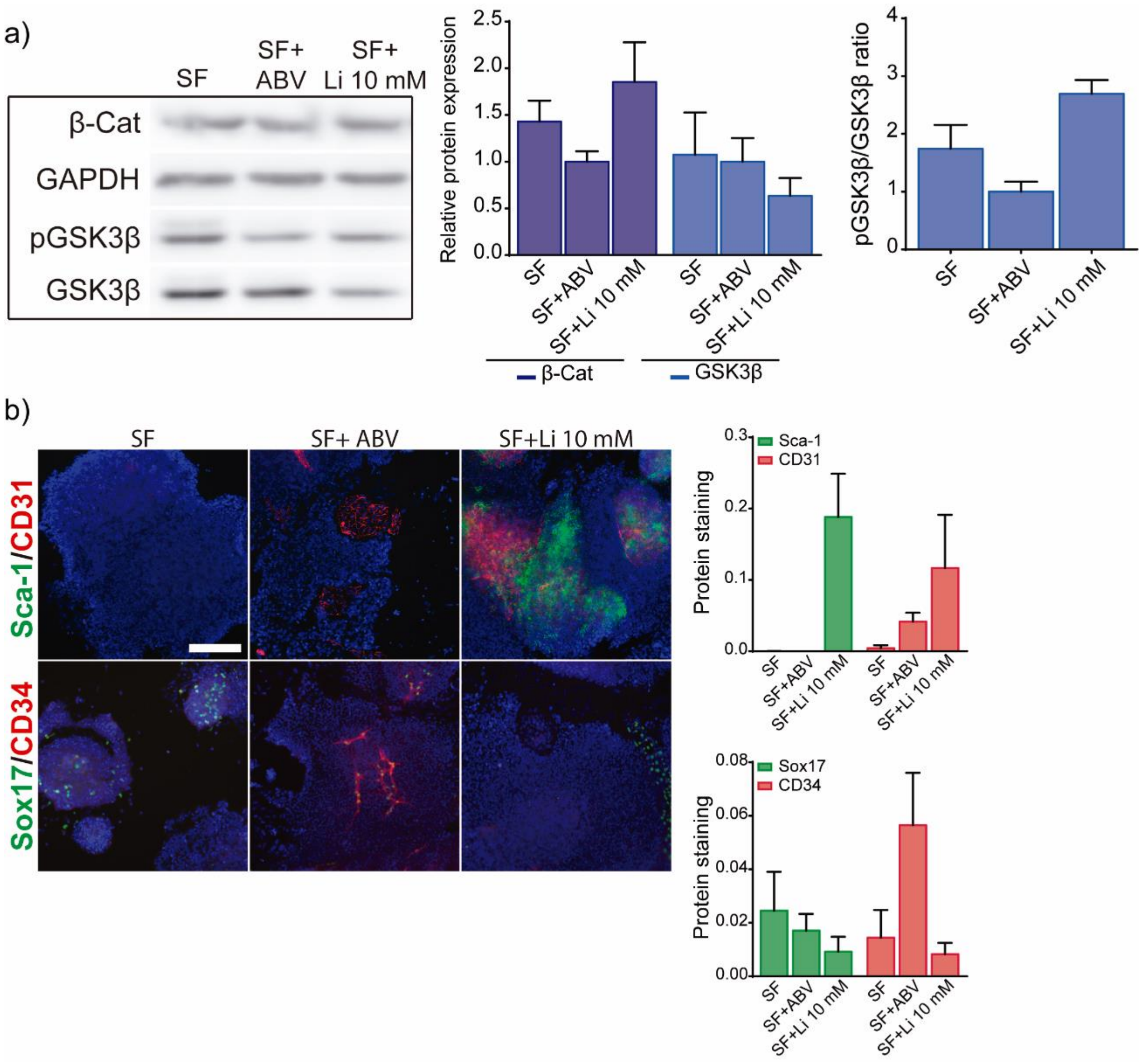


**Supplementary Figure 6**. ESCs treated with 10 mM $Li^{+}$ differentiated into Sca-1 and CD31-+ cells even in serum free (SF) differentiation medium.

ESCs were cultured in suspension in SF and SF supplemented with the combination of either Activin A, BMP4, VEGF or just 10 mM $Li^{+}$. After 4 days, spheroids were collected, disaggregated with trypsin and seeded in monolayer culture. Cells were then cultured for an additional 2 days to analyze:

a) Western blot analysis of GSK3β phosphorylation (S9), expression of GSK3β and β-catenin. GAPDH was used as loading control protein (n = 4).

b) Immunofluorescence analysis of markers related to endoderm (Sox17), endothelial lineage (CD34) and hemogenic endothelial lineage (Sca-1 and CD31). Scale bar: 200 µm.

Graphs show mean ± standard deviation. Significant differences were determined by ANOVA test; $*p < 0.05$.

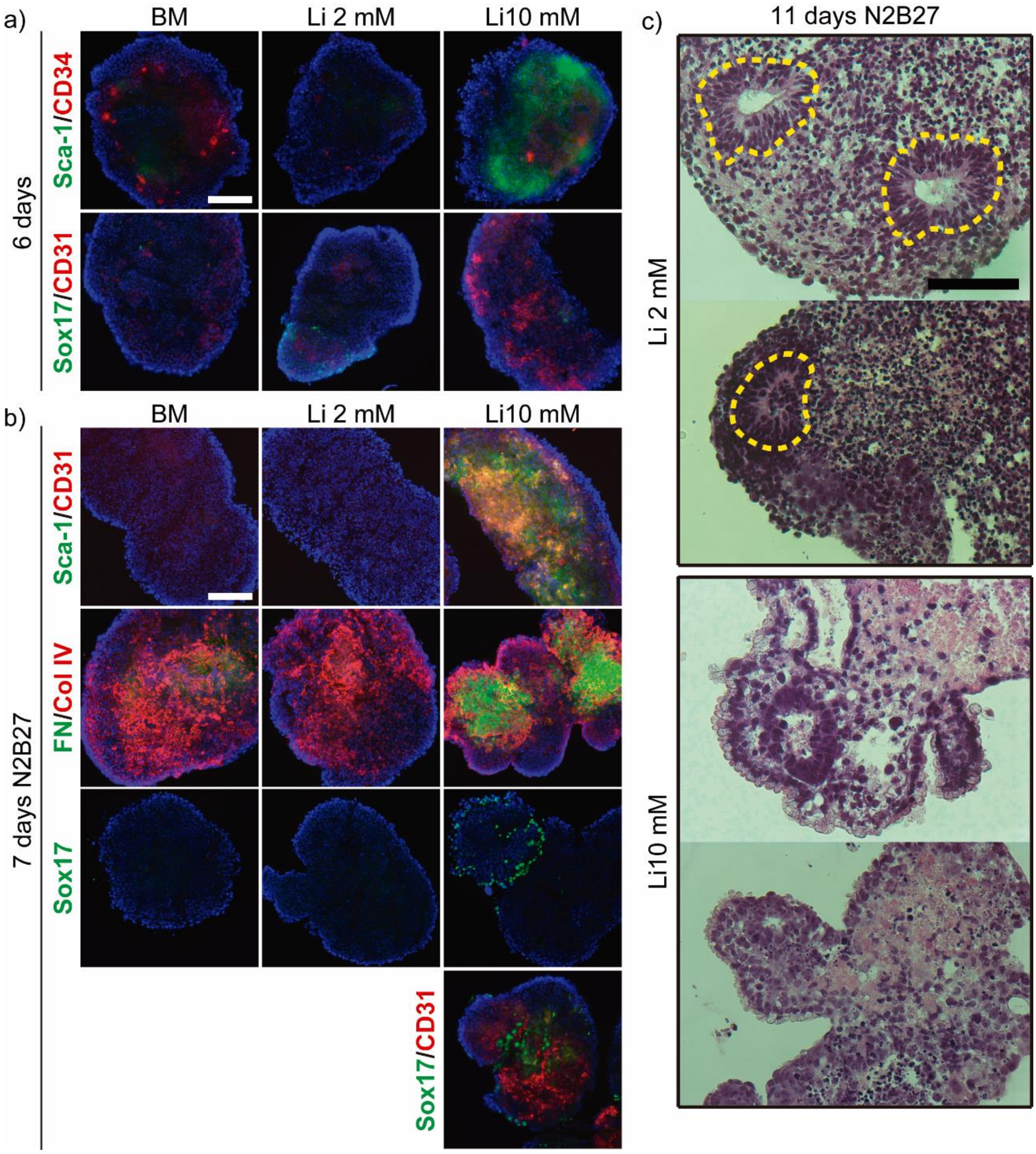


**Supplementary Figure 7**. ESCs treated with 10 mM $Li^{+}$ differentiate into hemogenic endothelial cells even when cultured in suspension.

a) Immunofluorescence analysis of cryosections (10 µm) of spheroids developed from ESCs cultured for 6 days in BM and BM supplemented with different concentrations of $Li^{+}$ (Scale bar: 100 µm). ESCs were seeded at 150,000 cells/ml in non-treated 6-well plates to develop the spheroids. The expression of mesoderm markers (Sca-1, CD34 and CD31) and endoderm marker (Sox17) were analyzed in order to compare differentiation dynamics with monolayer culture.

b) Immunofluorescence analysis of cryosections (10 µm) of spheroids previously developed for 6 days and then maturated for an additional 7 days in maturation medium N2B27 (Scale bar: 100 µm). The expression of hematopoietic stem cell precursor markers (Sca-1, CD31 and Sox17) and proteins related to the extracellular matrix of endothelial cells (fibronectin (FN) and collagen IV (Col IV)) were analyzed.

c) Hematoxylin/Eosin staining of histological sections (6 µm) of the spheroids developed from ESCs cultured with 2 and 10 mM $Li^+$ and then maturated with N2B27 for 11 days (Scale bar: 100 µm). The area inside dashed lines (yellow) highlight neural rosettes related to neural differentiation in EBs generated in presence of 2 mM $Li^+$. This was not observed for the spheroids in the presence of 10 mM $Li^+$.

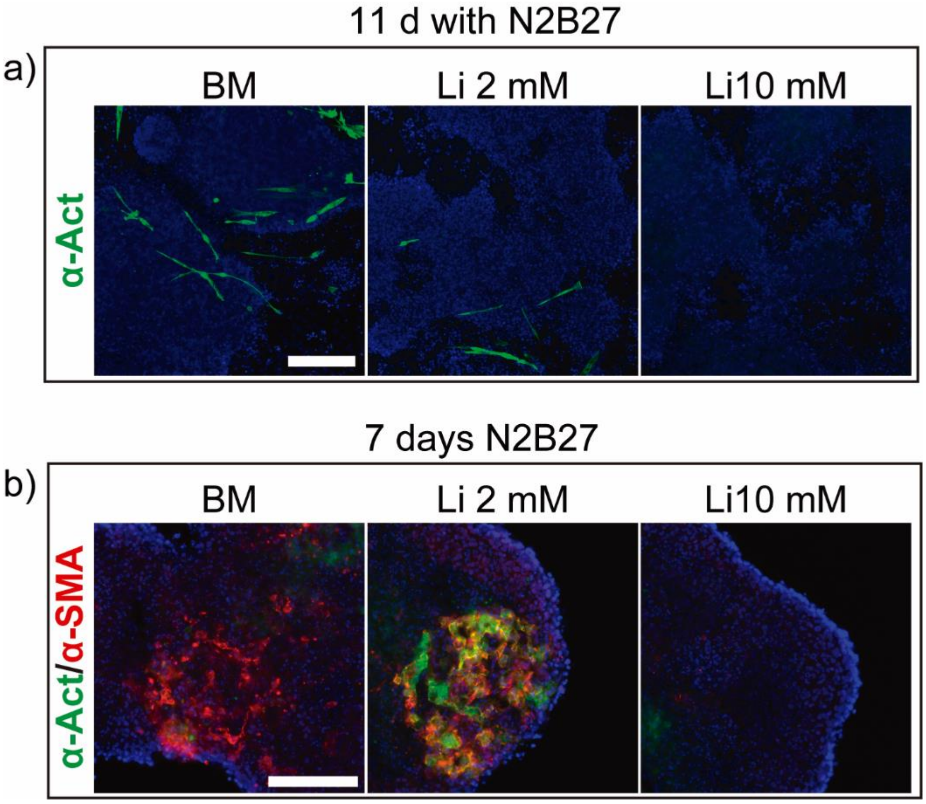


**Supplementary Figure 8**. ESCs treated with 10 mM $Li^+$ do not show cardiac differentiation features.

a) Immunofluorescence detection of markers related to cardiac lineage (sarcomeric α-actinin) in ESCs pre-cultured in BM and the presence of 2 and 10 mM $Li^+$ and then maturated with N2B27 for 5 days and 11 days (Scale bar: 200 µm).

b) Immunofluorescence analysis of cryosections (10 µm) of spheroids previously developed for 6 days and then maturated for an additional 7 days in maturation medium N2B27 (Scale bar: 100 µm). The expression of cardiac lineage markers was analyzed (sarcomeric α-actinin and α-smooth muscle actin). Scale bar: 100 µm.

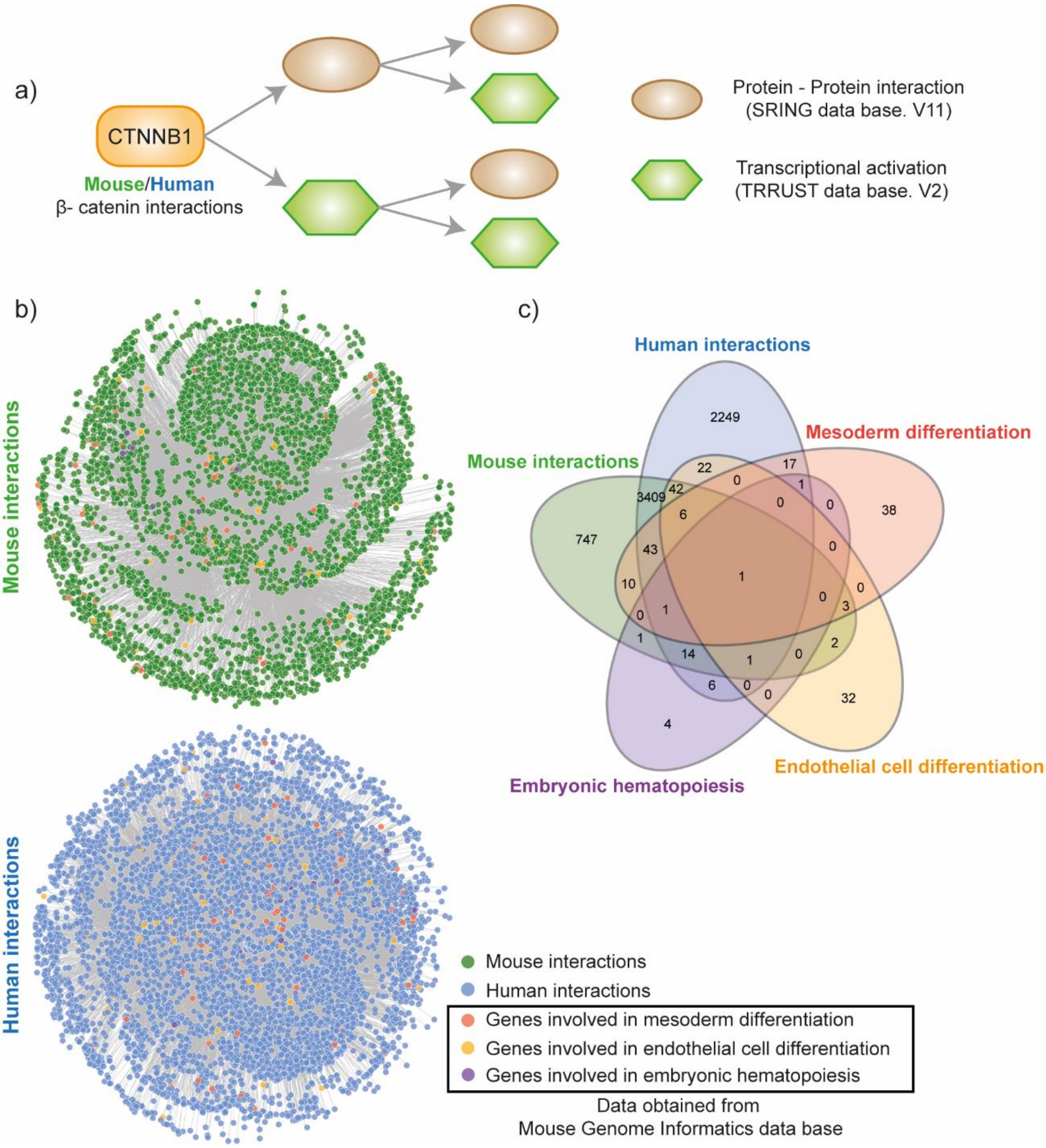


**Supplementary Figure 9**. Set of genes activated downstream P53 (Trp53) and β-catenin (Ctnnb1) and their implication in the differentiation of ESCs into definitive HSCs progenitors. Data was obtained from TRRUST database of human and mouse transcriptional regulatory networks. HE: Hemogenic endothelium, HSC: hematopoietic stem cells.
